# Combining Stem Cell Rejuvenation and Senescence Targeting to Synergistically Extend Lifespan

**DOI:** 10.1101/2022.04.21.488994

**Authors:** Prameet Kaur, Agimaa Otgonbaatar, Anupriya Ramamoorthy, Ellora Hui Zhen Chua, Nathan Harmston, Jan Gruber, Nicholas S. Tolwinski

**Affiliations:** Division of Science, Yale-NUS College, Singapore 138527, Singapore; Program in Cancer and Stem Cell Biology, Duke-NUS Medical School, Singapore 169857, Singapore; Department of Biochemistry, NUS, Singapore

## Abstract

Why biological age is a major risk factor for many of the most important human diseases remains mysterious. We know that as organisms age, stem cell pools are exhausted while senescent cells progressively accumulate. Independently, induction of pluripotency *via* expression of Yamanaka factors (*Oct4, Klf4, Sox2, c-Myc;* OKSM) and clearance of senescent cells have each been shown to ameliorate cellular and physiological aspects of aging, suggesting that both processes are drivers of organismal aging. However, stem cell exhaustion and cellular senescence likely interact in the etiology and progression of age-dependent diseases because both undermine tissue and organ homeostasis in different if not complementary ways. Here, we combine transient cellular reprogramming (stem cell rejuvenation) with targeted removal of senescent cells to test the hypothesis that simultaneously targeting both cell-fate based aging mechanisms will maximize life and health span benefits. We show that these interventions protect the intestinal stem cell pool, lower inflammation, activate pro-stem cell signaling pathways, and synergistically improve health and lifespan. Our findings suggest that a combination therapy, simultaneously replacing lost stem cells and removing senescent cells, shows synergistic potential for anti-aging treatments. Our finding that transient expression of both is the most effective suggests that drug-based treatments in non-genetically tractable organisms will likely be the most translatable.

## INTRODUCTION

Life is a constant struggle. This is true at cellular and molecular levels where tissue homeostasis requires constant surveillance, repair and replacement of cells damaged or lost due to intrinsic and extrinsic insults (*1*). Stem cells play a pivotal role in this tissue homeostasis by providing a reservoir of pluripotent precursor cells, needed to replace fully differentiated cells that are lost or damaged (*2*). At the opposite end of the cell-fate spectrum are senescent cells, or cells that have permanently withdrawn from the cell cycle (*3*). Cellular senescence can be replicative, where it is triggered by telomere shortening or mediated by stochastic damage, such as oxidative damage to DNA. Senescent cells can also arise as a response to oncogene activation to oppose transformation and cancerous growth (*4*). By entering permanent replicative arrest, senescent cells prevent mutations from expanding, thereby providing a sink for genotoxic damage. However, the senescent state does not simply result in passive replicative arrest but instead leads to transcriptional changes causing resistance to apoptosis and increased secretion of pro-inflammatory signaling molecules, a process known as Senescence Associated Secretory Phenotype (SASP). Senescent cell induced SASP in turn promotes inflammation and contributes to age-dependent dysfunction and to the development of age-related diseases (*5*).

While the number of stem cells decreases in aging animals, senescent cells accumulate with age (*6*). Manipulating cell fates by cellular reprogramming (to rejuvenate somatic cells) and by senolytic interventions (to remove senescent cells) are two promising approaches to restore homeostasis in aged individuals and to prevent age-dependent diseases. Cellular reprogramming allows differentiated cells to regain plasticity and to take on more stem cell-like qualities. A major step towards this goal was the demonstration of cellular reprogramming of terminally differentiated cells into pluripotent embryonic-like stem cell states (*7*). Importantly, such reprogramming reverses epigenetic aging marks, demonstrating that even mature, terminally differentiated cells can be returned to a younger state (*8*). While continuous expression of the Yamanaka factors (*Oct4, Klf4, Sox2, c-Myc*; OKSM) in mice led to the formation of teratomas and decreased lifespan (*9, 10*), repeated short term expression in adult mice succeeded in ameliorating cellular and physiological signs of aging (*11-13*). Subsequently, several studies have suggested that this approach can be applied to human aging and age-related disease (*14-18*), and cycling expression can rejuvenate stem cells in vitro (*19*).

Ablation of senescent cells has been shown to reverse tissue dysfunction and extend healthspan in mice (*20, 21*). A recent study using a senolytic construct (FOXO4-DRI peptide) that induced apoptosis in senescent cells, by interfering with the binding of p53 to FOXO4 thereby freeing p53 to activate apoptosis, showed that the clearing of senescent cells both counteracted senescent cell induced chemotoxicity and restored age-dependent declines in physical performance, fur density, and renal function in aging mice (*22*). Several studies have further explored applications of different senolytic strategies to ameliorate age-related decline and disease (*6, 23-26*).

Accumulation of senescent cells and loss of stem cells are not independent processes. Through SASP, senescent cells release large amounts of pro-inflammatory cytokines which contribute to chronic inflammation and mTOR activation, ultimately leading to stem cell exhaustion (*27*). This interaction suggests that senolytic therapies might interact with cellular reprogramming strategies in delaying age-dependent decline and disease. We have previously explored drug-drug interactions as synergistic aging interventions (*28*), and here we ask whether a combinatorial treatment of OKSM and senolytic (Sen) expression could mitigate or reverse the effects of aging more efficiently than either intervention alone. To test this hypothesis, we induced expression of OKSM, Sen and an OKSM-Sen combination in adult flies and compared their effects on health and lifespan. We find that each treatment alone had limited benefits, with OKSM alone benefiting maximum lifespan at the expense of healthspan while Sen expression alone increased mean lifespan but had no effect on maximum lifespan. In contrast, animals subjected to the combined intervention experienced substantially longer mean and maximum lifespan. Our data is consistent with a synergistic interaction between the two interventions, simultaneously rejuvenating stem cells and removing senescent cells.

## RESULTS

To test the interaction between senolytic removal of senescent cells and cellular reprograming, we designed a model combining these two interventions in an inducible overexpression system in *Drosophila*. First, we used the four Yamanaka factor based OKSM approach as this had been previously shown to induce pluripotent stem cells in mice (*7*), humans (*29-31*) and non-mammalian vertebrate and invertebrate species (*32*). To make a senolytic factor for *Drosophila*, we took advantage of the mouse sequence (FOXO4-DRI (*22*)) to design an orthologous peptide optimized to disrupt the *Drosophila* p53-dFoxO interaction. We then characterized effects of these two interventions independently as well as in combination.

We began by looking at the effect of OKSM and Sen on stem cells in an intestinal stem cell (ISC) model (*33, 34*). We chose to investigate phenotypic effects specifically in the digestive system of *Drosophila* (S. Fig. 1). As in mammals, the *Drosophila* gastric lining has a high turnover of cells which is enabled by stem cell pools that replenish the epithelia (*33*). Age-dependent loss of stem cells and degradation of barrier function has been shown to contribute to age-dependent functional decline and mortality in *Drosophila* (*35*). The *Drosophila* gut is composed of four cell types: enterocytes (ECs or absorptive cells), enteroendocrine (EEs or secretory cells), enteroblasts (EBs or transit amplifying cells) and intestinal stem cells (ISCs). ISCs rest on the external surface of the gut epithelium away from the gut lumen, and divide symmetrically to generate more ISCs, or asymmetrically to form EBs (*36*). The small, bright green cells or ISCs, can be observed either by expression of the stem cell determinant *escargot* (esgGal4>UAS-GFP), or by using a marker of Wnt activation, β-catenin (*armadillo* or *arm* in *Drosophila*), observable by GFP construct inserted into the endogenous locus (*37*).

We looked at the effect of expression of the two factors separately or together over a time course of 28 days. We observed a marked increase in ISC numbers starting at day 7 and continuing into day 28 in all three experimental conditions (Fig. 1). We observed an increase in ISCs and transit amplifying cells in OKSM expressing epithelia (Fig. 1B, F, J, N), an effect likely explained by the presence of Myc and suggesting that stem cell exhaustion may occur (*38*). We saw a similar increase in ISCs and transit amplifying cells in Sen expressing epithelia possibly due to the effect of p53 on stem cells (*39*). The increase in ISC numbers in animals expressing OKSM was expected, but surprisingly we observed a large increase in ISCs when Sen was expressed (Fig. 1C). Overall, the treatments showed higher numbers of stem cells over time as compared to wildtype flies. We next looked at lifespan effects. Continuous expression of OKSM is detrimental in mice while repeated short-term expression was beneficial (Ocampo et al., 2016). We expressed OKSM in ISCs which led to a significant detriment in lifespan (S. Fig. 2A).

**Figure 1:**
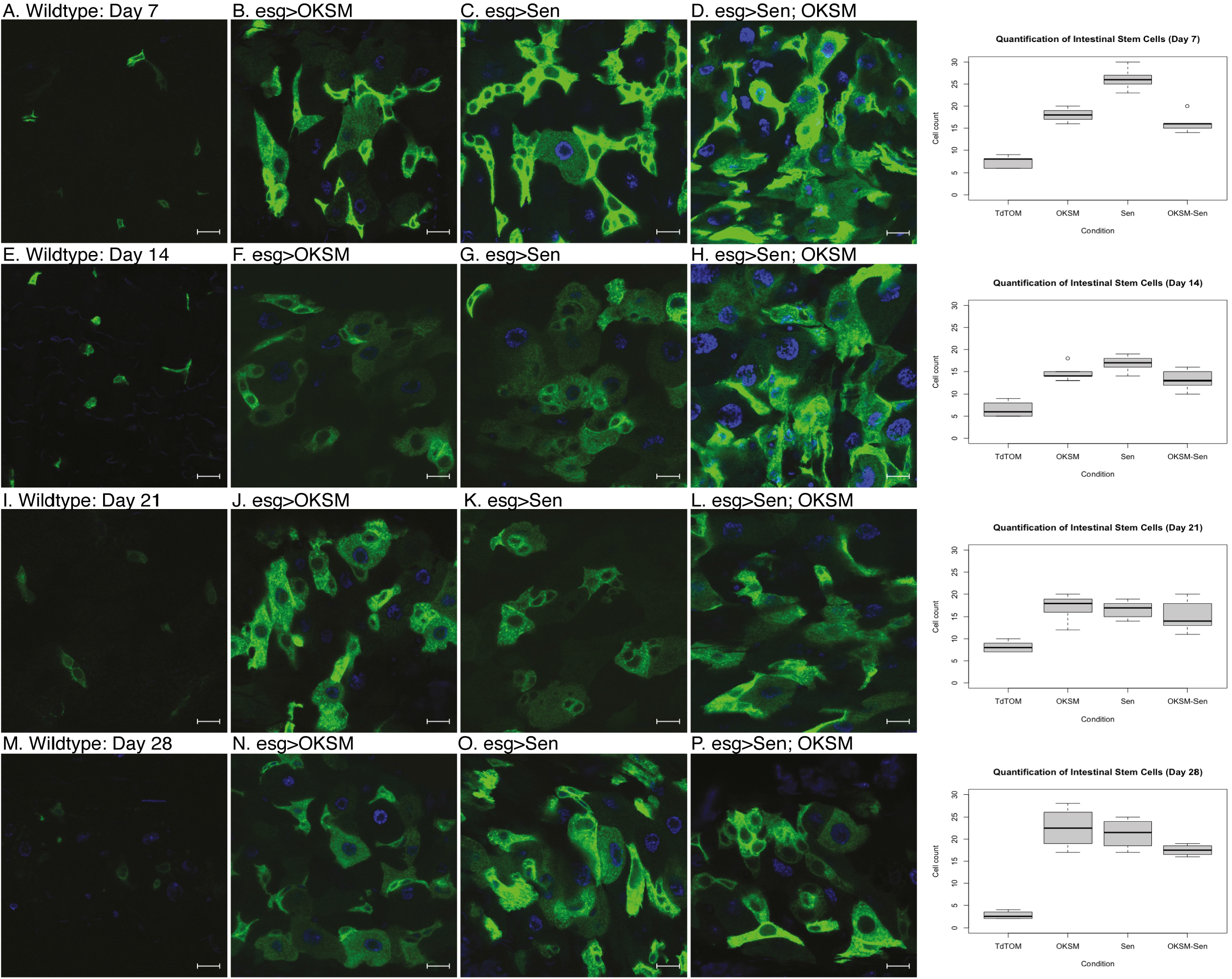
Expression of OKSM, Sen and OKSM-Sen led to increased stem cell proliferation over time. **(A)** esgGal4, UAS-GFP; tubGal80^ts^> UAS-TdTomato control flies show a small number of stem cells (n<10) and few enteroblasts during after seven days. Expression of OKSM **(B)**, Sen **(C)** or both Sen and OKSM **(D)** led to an increase in both ISCs and EBs with the highest number of ISCs observed in the Sen condition as quantified (Right). On day 14, little change was observed in control files **(E)**, but consistently higher numbers of ISCs were maintained in all three experimental conditions **(F-H, quantified Right)**. Day 21 showed little change from day 14 with similar numbers of ISCs observed in the control flies **(I)** and consistently higher numbers in the three experimental conditions **(J-L, quantified Right)**. By day 28, the number of ISCs was markedly decreased in control flies **(M)**, while all three experimental conditions maintained ISC numbers **(N-P, quantified Right)**.

To look more closely at the effect of Sen and OKSM, we determined the transcriptional profiles of the three conditions as compared to wildtype. We dissected the midguts of flies expressing OKSM, Sen, OKSM and Sen (OKSM-Sen), and control flies (WT) expressing only a fluorescent protein, either in the ubiquitous expression (*i*.*e*., under the control an *armadillo* driver) or the ISC-restricted expression model (*i*.*e*., under control of an *escargot* driver) and performed RNA-seq (Fig. 2, S. Fig. 3).

**Figure 2:**
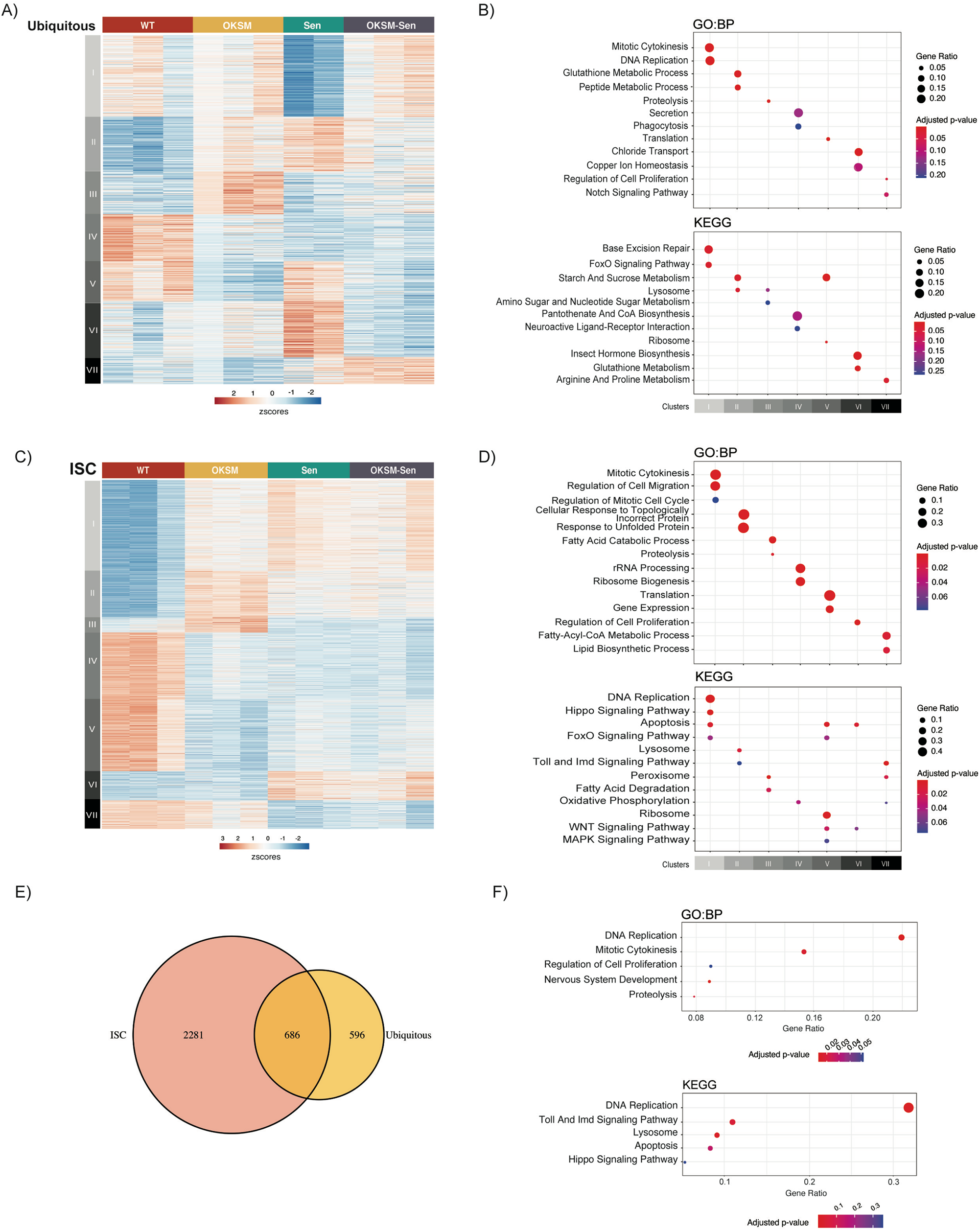
Gene expression changes in the *Drosophila* gut in Sen, OKSM and Combination treatment. **(A)** Heatmap of gene expression in ubiquitous expression experiments with armGal4; tubGal80^ts^ > UAS-TdTomato (WT), armGal4; tubGal80^ts^ > UAS-OKSM (OKSM), armGal4; tubGal80^ts^ > UAS-Sen (Sen) and armGal4; tubGal80^ts^ > UAS-Sen; UAS-OKSM (OKSM-Sen) dissected midguts showing seven clusters with differing expression patterns across the four conditions. (**B)** Gene Ontology and KEGG Pathway Enrichment analysis of ubiquitous expression highlighting key signaling and metabolic pathways associated with the individual clusters (FDR < 10%). **(C)** Heatmap of gene expression changes in stem cell only expression experiments with esgGal4; tubGal80^ts^ > UAS-TdTomato (WT), esgGal4; tubGal80^ts^ > UAS-OKSM (OKSM), esgGal4; tubGal80^ts^ > UAS-Sen (Sen) and esgGal4; tubGal80^ts^ > UAS-Sen; UAS-OKSM (OKSM-Sen) dissected midguts again showing seven clusters with differing expression patterns across the four conditions. **(D)** Gene Ontology and KEGG Pathway Enrichment analysis of ISC only highlighting key signaling and metabolic pathways associated with the individual clusters (FDR < 10%). **(E)** Venn diagram showing overlap in genes from ubiquitous and ISC-restricted expression. **(F)** Gene Ontology and KEGG Pathway Enrichment analysis of the major pathways affected in both models.

In the ubiquitous expression model, 1282 genes (FDR < 10%) were identified as significantly differentially expressed (Table S1). Clustering of these genes identified seven distinct clusters, each representing groups of genes with similar expression profiles across the four conditions (Fig. 2A). Each of the seven clusters was enriched for distinct pathways and processes (Fig. 2B, Supplementary Table 1). Cluster I (N = 300) contained genes that were downregulated by Sen but were unaffected in the other conditions. Genes in this cluster were associated with cytokinesis, cell cycle and DNA replication. These would be expected results as p53 release from FoxO should lead to apoptosis rather than cell proliferation. Genes in Cluster II (N = 201) were upregulated in all conditions compared to WT. This cluster was enriched for processes associated with Glutathione and sugar metabolism and lysosomal activity, which are related to tissue building and repair. Cluster III (N = 157) consisted of genes that were upregulated only in the OKSM condition and was enriched for lipid and amino acid metabolism. Cluster IV (N = 173) consisted of genes that were downregulated by all conditions relative to WT and was enriched for genes associated with secretion and phagocytosis. Cluster V (N = 147) represented genes that were downregulated in both OKSM and in the combined OKSM-Sen conditions and contained genes related to translation. Cluster VI (N = 204) contained genes that were upregulated specifically by Sen alone and consisted mainly of genes involved in homeostasis. Cluster VII (N = 100) contained genes upregulated by the OKSM-Sen combination which were involved in Arginine and Proline amino acid metabolism. Overall, our results suggested that genes affected by expression of Sen correlated with lower cell division and FoxO signaling, and a higher level of amino acid metabolism. Expression of OKSM upregulated amino acid and lipid metabolism and proteolysis while downregulating translation and sugar metabolism.

In the ISC-restricted expression model, we identified 2967 genes (FDR < 10%) as significantly differentially expressed across conditions (Supplementary Table 2), which were subsequently clustered into seven distinct groups (Fig. 2C-D, Fig. 3). Cluster I (N = 773) contained genes that were upregulated in all three conditions relative to WT. Genes in this cluster were associated with cytokinesis, cell cycle, cell migration and DNA replication, further supporting that in ISCs expression of these factors, in any combination, results in increased proliferation and migration of ISCs. Genes in Cluster II (N = 397) were upregulated in OKSM-expressing ISCs. This cluster was enriched for processes associated with the misfolded protein response, Toll signaling and the lysosome. Cluster III (N = 128) consisted of genes that were specifically upregulated by OKSM, and downregulated in the other conditions, and was enriched for fatty acid degradation and peroxisome function. Cluster IV (N = 566) consisted of genes that were downregulated in all conditions relative to WT. Cluster IV was enriched for genes involved in ribosome biogenesis, fatty acid biosynthesis and oxidative phosphorylation. Cluster V (N = 613) contained genes that were downregulated the most in OKSM, but also in the other conditions studied, and contained genes related to protein translation and signaling pathways. Genes in Cluster VI (N = 248) were specifically upregulated following expression of Sen or OKSM-Sen, and contained genes involved in major developmental signaling pathways and apoptosis. Cluster VII (N = 242) contained genes downregulated by Sen and the OKSM-Sen combination, and was enriched for genes involved in Toll and mTOR signaling and various metabolic and homeostatic processes. Overall, our transcriptional analysis suggested that gene expression was affected by the expression of Sen or OKSM individually, but that there was not a distinct group that was differentially expressed specifically in response to the combination (Sen-OKSM). Overall, OKSM expression upregulated misfolded protein response and Toll signaling, while downregulating Insulin secretion and protein translation, whereas Sen expression activated Wnt and Hedgehog signaling and downregulated Toll and mTOR pathways. Taken together, this indicates that the observed changes in mean and maximum lifespan are associated with two distinct transcriptional programs and is not the result of Sen-OKSM specifically activating or repressing a shared transcriptional pathway.

**Figure 3:**
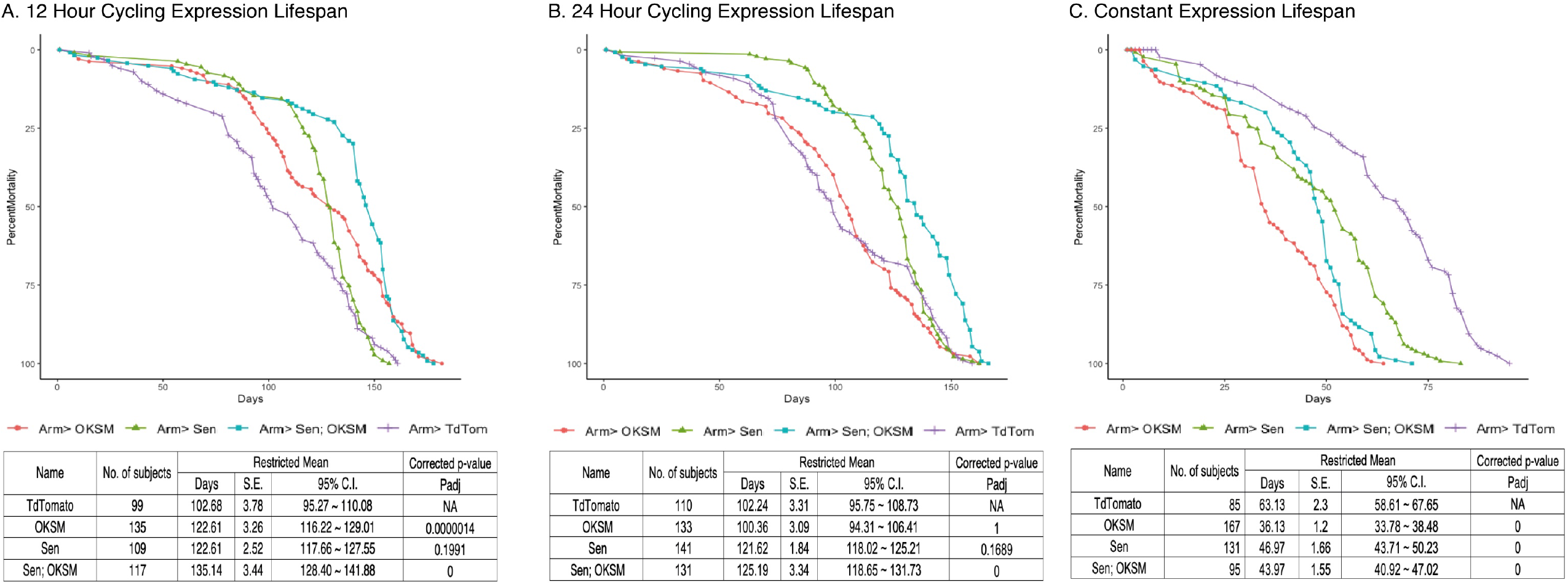
Cycling OKSM and Sen expression leads to lifespan and health span extension while combined OKSM-Sen expression increases both. **(A)** Survival curve for armGal4; tubGal80^ts^ > UAS-TdTomato (TdTom), armGal4; tubGal80^ts^ > UAS-OKSM (OKSM), armGal4; tubGal80^ts^ > UAS-Sen (Sen) and armGal4; tubGal80^ts^ > UAS-Sen; UAS-OKSM (Sen; OKSM) where expression is limited to one twelve-hour period twice per week through a temperature shift. At 25º the temperature sensitive gal80 protein ceases to inhibit gal4 from driving expression from UAS enhancers, allowing for a targeted expression window when flies were shifted from 18º to 25º. Expression of OKSM, Sen and Sen; OKSM in adult female flies resulted in increased lifespan as compared to control flies (TdTom). Mean and maximum lifespans are shown along with corrected p-values. **(B)** Survival curve for the same experiment but with 24 hours of expression twice per week induced by a temperature shift of 18º to 25º. There were similar benefits of expression but reduced compared to the 12-hour expression experiment. Mean and maximum lifespans are shown along with corrected p-values. **(C)** Survival curve for flies expressing OKSM, Sen and Sen; OKSM but maintained at 25º throughout their lifespans. The overall lifespans are shorter due to the higher temperature, but in addition all three experimental conditions show detriments to both mean and maximum lifespans. Mean and maximum lifespans are shown along with corrected p-values. A P-value of 0 reflects P < 1.0 * 10^−10^.

Next, we compared the sets of differentially expressed genes identified in both models to determine if there was also a shared core transcriptional program was altered in both systems. Overall, 686 genes were differentially expressed in both models (Fig 2E, p<1×10^−6^), with these genes being enriched for DNA replication, regulation of epithelial cell migration, mitosis, inflammation, various metabolic processes and specific developmental signaling pathways (Fig. 2F). The majority of these genes have similar responses to transgene expression in either model, while some exhibit differences in their response between models.

To evaluate lifespan effects under optimized conditions, we designed two approaches for cycling expression to overcome the continuous expression detriment. We first used a drug induced expression model where the polycistronic OKSM transgene, under the transcriptional control of UAS regulatory sequences, was driven by the Actin-Switch-GAL4 driver activated by RU486 (*40*). Flies were placed on fresh food supplemented with the drug weekly leading to periodic, ubiquitous expression. We found that OKSM expression alone resulted in mean lifespan extension in both male and female flies, with female flies showing an increase maximum lifespan as well (S. Fig. 2B). The advantage of this system was that flies could be cultured at higher temperatures reducing the overall length of lifespan studies and allowing rapid confirmation of lifespan benefits. However, this system does not allow for more precise control of expression due to drug half-life, consumption, and distribution to all tissues. For this we turned to a temperature sensitive expression system where a ubiquitous GAL4 driver was combined with a ubiquitous, temperature sensitive GAL80 inhibitor. This system allowed us to generate adults with no embryonic expression, and by modulating the temperature of culture, we were able to precisely induce expression in all tissues for defined periods ranging from constant to once per week. We found that continuous expression of OKSM was detrimental (Fig. 3C), expression for 24 hours twice per week was mildly beneficial (Fig. 3B), and for 12 hours twice per week showed lifespan extension (Fig. 3A).

We carried out similar optimization experiments for the senolytic peptide (Sen) alone and found that, in terms of median lifespan, continuous expression was also detrimental while expression for either 24 or 12 hours twice per week resulted in significant lifespan extension (Fig. 3). Having established conditions under which each individual intervention was beneficial, we then tested whether simultaneous removal of senescent cells (Sen) and cellular reprogramming (OKSM) would result in additive or synergistic benefits in aging flies. The combined intervention again was detrimental when expression of Sen and OKSM was induced continuously, but extended both maximum lifespan and median lifespan when expressed for 24 hours once per week (Fig. 3). Most striking was the significant mean and maximum lifespan extension noted in flies with OKSM and Sen expressed together for 12 hours twice per week (Fig. 3A).

We reasoned that longer-lived flies should maintain stem cell pools for longer due to stem cell rejuvenation through OKSM expression. To test this hypothesis, we examined the midguts from flies expressing the various transgenes over a time course of 28 days. We visualized ISCs in gastric epithelia of flies with cycling expression of OKSM, Sen or both at four weeks (Fig. 4A-D), eight weeks (Fig. 4G-J), and twelve weeks (Fig. 4M-P). We observed and quantified a higher number of ISCs in flies when OKSM was expressed (Fig. 4E-F, K-L, Q-R).

**Figure 4:**
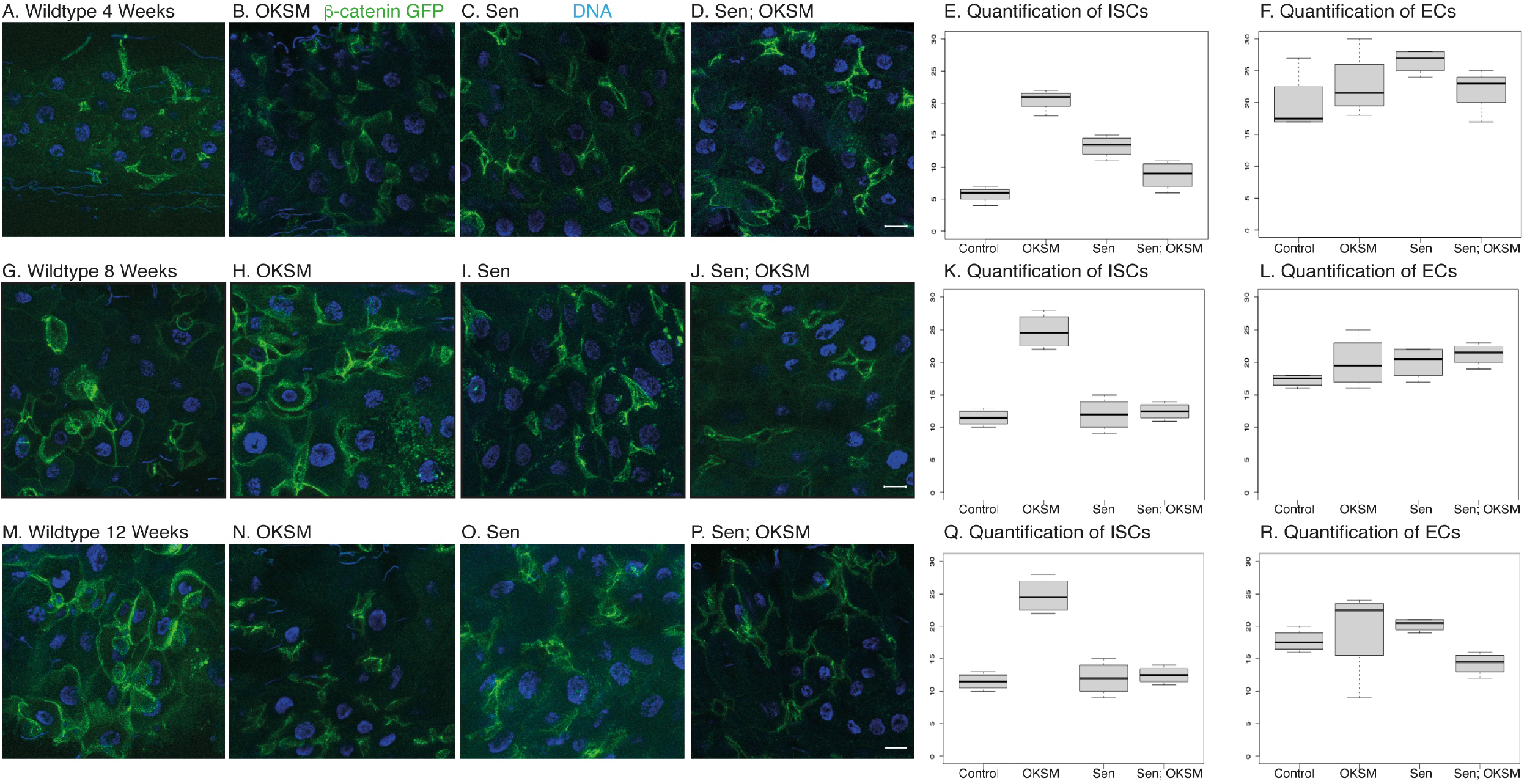
Cycling OKSM expression maintains a larger pool of intestinal stem cells over time. **(A)** We visualized stem cells with b-catenin-GFP (arm^GFSTF^) which is enriched in ISCs due to Wnt signaling. Flies were cycling expression through temperature shifts from 18º to 25º for twelve hours once a week. arm^GFSTF^; armGal4; tubGal80^ts^ > UAS-TdTomato control flies show a small number of stem cells and few enteroblasts after four weeks. Expression of OKSM **(B)**, Sen **(C)** or both Sen and OKSM **(D)** led to an increase in both ISCs with the highest number of ISCs observed in the OKSM condition as quantified **(E-F)**. After eight weeks, the number of ISCs in the OKSM condition was much higher in the OKSM **(H)** flies as compared to wildtype **(G)** and Sen **(I)** or Sen; OKSM **(J)** quantified **(K-L)**. At twelve weeks, OKSM **(N)** flies maintain high numbers of stem cells while the other conditions **(M, O-P)** show fewer as quantified **(Q-R)**.

Our observations for periodic expression of Sen were consistent with data from mice subjected to senolytic interventions. Sen flies experience a substantial increase in mean but not in maximum lifespan, indicating compression of mortality with excess late deaths compensating for protective effects earlier in life. The same is not true for OKSM flies which experience a statistically significant maximum lifespan extension with both 24h and 12h induction. Strikingly, simultaneous application of Sen and OKSM, especially for 12h induction, result in a mortality trajectory that combines beneficial features of both individual interventions and result in mean and maximum lifespan extension benefit that exceed either. To further investigate this interaction, we performed a quantitative analysis of age-dependent mortality.

Biological aging is defined by an exponential increase in mortality rate over time. Mathematically, this is expressed by the Gompertz–Makeham law of mortality (*41*):

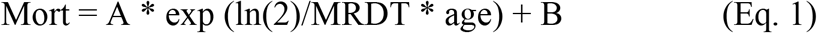

Where A is the initial mortality rate in young animals, B is the age-independent mortality and MRDT is a characteristic time interval over which age-dependent mortality doubles. To quantify the impact of single interventions and the combined intervention on age-dependent mortality, we followed an approach recently described by Axel Kowald and Tom Kirkwood, fitting Gompertz– Makeham survival functions to our experimental survival data (*42*). This fit yielded estimates for the initial mortality A and the MRDT parameters of flies (Supplementary Table 3). All fits resulted in a good agreement between experimental data and the Gompertz curve, with a mean residual standard error of 0.03 (3% survival) across all conditions (Fig. 5 B, D, F, Supplementary Table 3). Mortality trajectories were then visualized by plotting the logarithm of mortality against age (Fig. 5 A, C, E). In this graph the initial mortality A is the intercept of the mortality trajectory at time zero while the slope of the line is proportional to the inverse of the MRDT. Sen-driven lifespan extension showed a substantial decrease in early mortality A, relative to control. However, this decrease was associated with a significant penalty in the form of age-acceleration (decreased MRDT). For the 12h induction, early mortality decreases almost 100-fold while MRDT decreases from 22.7 days to 7.9 days (p<0.05). In other words, while initial mortality is substantially lower following Sen treatment, the treated flies age approximately 290% faster than WT. This pattern was consistent with previously described mouse data and explains why Sen treatment results in mortality compression with increased mean but not maximum lifespan. By contrast, the impact of OKSM induction on initial mortality and MRDT was much smaller. OKSM induction for 24h and 12h significantly reduced initial mortality A by 46% and 30.1%, (p<0.05), respectively (Supplementary Table 3). While OKSM was also associated with a slight age acceleration penalty in terms of MRTD, this effect was much smaller than for Sen. For 12h OKSM induction, MRDT only decreased from 22.7 days to 19.6 days (p<0.05); a 15.8% increase in aging rate. The result of this change can be seen when plotting log mortality as a function of age (Fig. 5 C, E). OKSM mortality was shifted downwards relative to control but ran largely parallel to control mortality, meaning that age-dependent mortality for 12h OKSM animals remained lower than for WT at later ages. This pattern explains why OKSM impacted maximum lifespan and was not associated with mortality compression.

**Figure 5:**
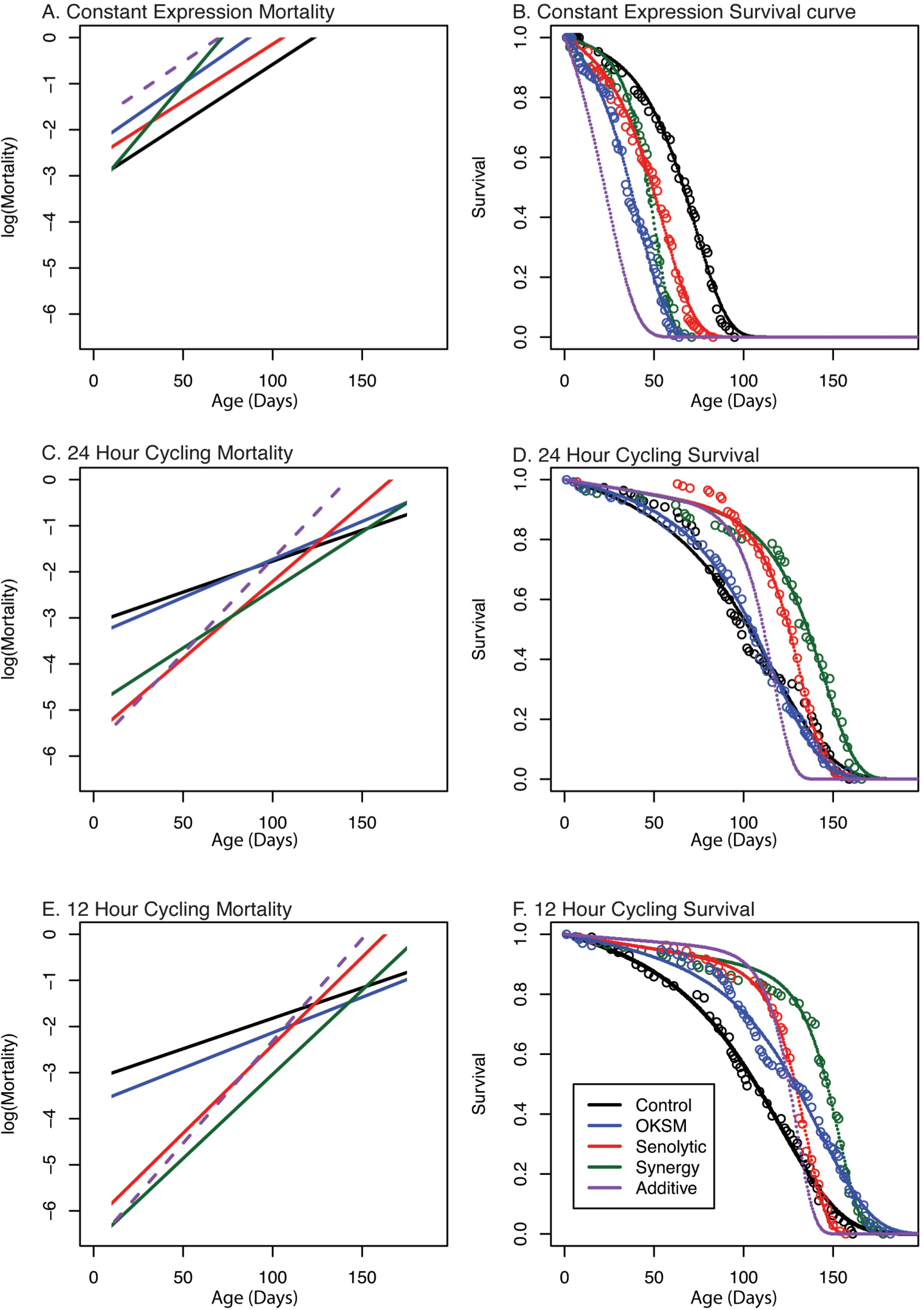
Gompertz–Makeham mortality and survival analysis demonstrates decreased early mortality with compensating increased aging rate in Sen and OKSM interventions. Survival data for each intervention are shown together with the best-fit survival curve on the right and the corresponding mortality trajectory are shown on the left. Initial log(Mortality) parameter A can be read as the intersection between mortality curve and y-axis at age zero. The slope of the mortality curve is proportional to 1/MRDT. Dashed purple line illustrates hypothetical mortality trajectory assuming additivity of effects elicited by OKSM and Sen. (A,B) mortality trajectories and survival curve for cohorts with continuous induction of OKSM, Sen or OKSM+Sen and controls. (C,D) mortality trajectories and survival curve for cohorts with induction of OKSM, Sen or OKSM+Sen for 24h every three days and matched controls.. (E,F) mortality trajectories and survival curve for cohorts with induction of OKSM, Sen or OKSM+Sen for 12h every three days and matched controls. Note that flies subject to continuous induction (Panels A and B) are permanently kept at 25º and are therefore aging more rapidly than flies cultured at 18º and induced for only for short periods. The slope of the mortality trajectory of controls in A is therefore approximately two times as larger, compared to that of controls in panels C and E. For exact MRTD and A parameter values and associated confidence intervals see: (Supplementary Table 2).

OKSM-Sen flies with 24h of induction experienced a significant reduction in the age-acceleration penalty (p<0.05) relative to 24h Sen-only flies (Fig. 5 C, Supplementary Table 3). OKSM-Sen flies with 12h induction also showed this trend, but the difference was not statistically significant (Fig. 5 E, Supplementary Table 3). In contrast, the initial mortality A was decreased by over 330-fold in 12h OKSM-Sen compared to WT. This means that initial mortality rate was significantly lower in OKSM-Sen than even in Sen only (p < 0.05), suggesting that adding OKSM to Sen partially rescued the age-acceleration penalty while further augmenting Sen benefits in terms of early mortality. The resulting survival trajectories consequently show both mean and maximum lifespan extension with mortality compression occurring only late in life. Mortality compression only becomes apparent after most WT animals have already died.

When investigating the interaction between Sen and OKSM treatments, it is useful to compare observed effects to a hypothetical survival and mortality trajectory constructed by assuming that the two interventions act independently (see materials and methods for details). Comparing the OKSM-Sen group to this hypothetical cohort (purple dashed lines in Fig. 5) revealed that these effects cannot be explained without a direct interaction between Sen and OKSM in terms of aging rate. On their own, both interventions accelerate aging rate (decrease MRDT) but improve early mortality. When combined, the intervention results in a reduction, rather than further increase, in the age-acceleration penalty while further augmenting early mortality benefits. This synergistic interaction between OKSM and Sen is the reason why the combined treatment improves both maximum and median lifespan more significantly than either of the two interventions alone. Mechanistically, these data imply a direct interaction between partial reprogramming *via* OKSM and the Sen-driven senescent cell apoptosis. Indeed, this is what we observed on the cellular level, as expression of Sen impacted the number of stem cells directly (Fig. 1) even without OKSM expression.

## DISCUSSION

Here we show that it is possible to extend both the mean and maximum lifespans by combining anti-aging strategies targeting two different ageing mechanisms related to cell fate. Pulsed expression of the four Yamanaka transcription factors to rejuvenate cells combined with a Senolytic factor kept flies healthier and extended their lives. Although not tested in our study, reprogramming leads to a change in DNA methylation and other epigenetic markers leading to a more youthful gene expression signature (*16*). The periodic removal of senescent cells leads to fewer chemotoxic molecules being produced and in rejuvenation of organs (*22*). Both interventions are rejuvenating in the sense that they reverse cellular tissue composition towards a more youthful state. The substantial reduction in initial mortality following both interventions is consistent with this mechanism. Here we report that these two interventions are more closely related than previously appreciated.

Senescent cells show persistent activation of the mTOR pathway, a state that promotes secretion of a wide range of signaling molecules, including proinflammatory cytokines (*43-45*). As these molecules are secreted, they have the potential to impact neighboring or distant cells increasing the number of senescent cells, impairing tissue homeostasis (*46, 47*). In the ubiquitous expression model, we found that expression of the Senolytic peptide led to a decrease in Tgfβ (−0.51 log_2_ Fold decrease, FDR=7.51×10^−2^) and the cytokine Upd3 (−1.0 log_2_ Fold decrease, FDR=2.91×10^−4^) compared to Control. Upd3 activates Jak/Stat signaling often related to stem cell activation upon injury leading to asymmetric divisions and lower numbers of stem cells (*48, 49*). Tgfβ is upstream of Upd3 and involved in promoting senescence (*50-52*). SASP-mediated activation of cytokines and mTOR therefore directly link age-dependent accumulation of senescent cells to accelerated loss of stem cells and declining capacity for repair and tissue regeneration. This mechanism suggests a model by which removal of senescent cells would promote increased resilience and improved maintenance of stem cell pools a phenotype we observe in the fly.

Although OKSM must function through partial reprogramming of cells, the exact mechanism of how this works in adult tissues is not entirely clear (*13, 16-18*). We observed changes in Hedgehog signaling recently proposed as a neuroprotective and life extending pathway (*53*) along with genes associated with cytokinesis and DNA replication. Importantly, we observe that OKSM has limited effect on maximum lifespan unless senescent cells are removed suggesting that SASP counteracts the benefits of rejuvenation. Previous studies have shown either OKSM or Sen to be anti-aging, but in both cases the effects did not affect maximum lifespan. In our combinatorial approach, we can now extend both mean and maximum. We have further established that both approaches can be studied in the much shorter-lived *Drosophila* model. We suggest that reprogramming accomplished through gene therapy, or another method combined with senolytic peptides or drugs could promote both tissue repair and reverse age-related decline.

## MATERIALS AND METHODS

### Molecular Cloning of Transgenes

The Oct4-2A-KLF4-2A-Sox2-IRES-Myc DNA fragment containing the human iPS factors was obtained from the OKSIM plasmid (OKSIM was a gift from Jose Cibelli, Addgene plasmid # 24603; http://n2t.net/addgene:24603; RRID:Addgene_24603)(*54*). Oct4-2A-KLF4-2A-Sox2 were amplified as one fragment with attB1 and att5r-flanked sites and recombined with the pDONR P1-P5r entry vector (Thermo Fisher Scientific). IRES-Myc was amplified with attB5 and attB2-flanked sites and recombined with the pDONR P5-P2 entry vector. MultiSite Gateway® Pro 2.0 recombination (Thermo Fisher Scientific) was used to recombine the 2 donor plasmids into the pUASg.attB.3XHA (A kind gift from J. Bischof and K. Basler, Zurich) (*55*) vector to obtain the OKSM gene cassette for expression in *Drosophila* (*56*).

The Senolytic (Sen) construct corresponded to amino acid 86 to 131 of the *Drosophila* Fork Head protein. The Sen construct was synthesized and transferred by Gateway cloning (Thermo Fisher Scientific) into pUASg.attB with C-terminal 3XHA tag (A kind gift from J. Bischof and K. Basler, Zurich) (*56*).

### Fly crosses and expression of constructs

For *Drosophila*, the transgenes were injected into attP2 (Strain#8622) P[CaryP]attP2 68A4 by BestGene Inc. (California) (*57*). Expression was driven by Actin-Switch-Gal4 (*40*), escargot-GAL4 (*33, 58*) and tubulin-GAL80^ts^ (*59*). All additional stocks were obtained from the Bloomington Drosophila Stock Center (NIH P40OD018537).

### Fly lines used in this study

Actin5C(−FRT)SwitchGAL4: BDSC 9431 (*40*)

UAS-Td-Tomato: BDSC 36328 (Joost Schulte and Katharine Sepp)

esg-Gal4, UAS-GFP; tub-Gal80^ts^, UAS-dCas9.VPR: BDSC 67069 (*33*)

arm^GFSTF^ MI08675-GFSTF: BDSC 60651 (*37, 60, 61*)

armGal4; tub-GAL80^ts^: BDSC 86327 (*59*) UAS-OKSM,

UAS-Sen (This study)

Fly crosses performed were:

1. Act5CGAL4-Switch x w; UAS-OKSM
2. ActGAL4-Switch x w; UAS-TdTomato
3. esg-Gal4, UAS-GFP; tubGal80^ts^ x w; UAS-OKSM
4. esg-Gal4, UAS-GFP; tubGal80^ts^ x w; UAS-Sen
5. esg-Gal4, UAS-GFP; tubGal80^ts^ x w; UAS-Sen; UAS-OKSM
6. esg-Gal4, UAS-GFP; tubGal80^ts^ x w; UAS-TdTomato
7. arm-Gal4, UAS-GFP; tubGal80^ts^ x w; UAS-OKSM
8. arm-Gal4, UAS-GFP; tubGal80^ts^ x w; UAS-Sen
9. arm-Gal4, UAS-GFP; tubGal80^ts^ x w; UAS-Sen; UAS-OKSM
10. arm-Gal4, UAS-GFP; tubGal80^ts^ x w; UAS-TdTomato
11. arm^GSFTF^; arm-Gal4; tubGal80^ts^ x w; UAS-OKSM
12. arm^GSFTF^; arm-Gal4; tubGal80^ts^ x w; UAS-Sen
13. arm^GSFTF^; arm-Gal4; tubGal80^ts^ x w; UAS-Sen; UAS-OKSM
14. arm^GSFTF^; arm-Gal4; tubGal80^ts^ x w; UAS-TdTomato

### Animal husbandry

*Drosophila* were maintained at standard humidity and temperature (25°C) with food containing 6g Bacto agar, 114g glucose, 56g cornmeal, 25g Brewer’s yeast and 20ml of 10% Nipagin in 1L final volume as previously described (*62*).

### Gut preparations

Adult fly midguts were dissected in 200 µl of 1x PBS in a PYREX™ Spot Plates concave glass dish (FisherScientific). The midguts were rinsed with PBS and stained with 10mg/ml Hoechst 33342 diluted 200 times in 1X PBS for 1min. Subsequently, the guts were carefully transferred onto a small droplet of 1x PBS on a 35mm glass bottom dish. Using fine forceps, the gut was repositioned to resemble its natural orientation. PBS was then removed from the area surrounding the gut, leaving a small amount of excess PBS to hold the gut in place and prevent desiccation. The 3mm glass bottom dish was then mounted onto the Zeiss LSM800 (Carl Zeiss AG, Germany) for imaging. For each construct, midguts from at least 3 flies were dissected and imaged at the 25% percentile from the anterior midgut (*63*).

### Fluorescence microscopy

Images were acquired on the Zeiss LSM 800 (Carl Zeiss, Germany) using the Plan-Apochromat 63X/1.4 Oil DIC M27 objective, 3% laser power for 488nm and 3% laser power for 405nm. Images were processed using the ZEN 2014 SP1 software (Carl Zeiss, Germany). Figures were made with Adobe Photoshop and Illustrator. Models were created with Biorender.com.

### Lifespan studies

For each experiment, more than 50 F_1_ flies were cultured at 25° or 18°C. Flies were counted daily noting the number of dead and censored subjects. Lifespans were scored every day. Flies that failed to respond to taps were scored as dead, and those that were stuck to the food were censored. Lifespan curves and statistical analysis of lifespan studies were performed using OASIS 2 (Online Application for Survival Analysis 2 (*64, 65*)). For studies using Actin-Switch, flies were moved to fresh vials with food supplemented by 200µM RU486 (mifepristone) weekly. For studies using the temperature sensitive expression inhibitor tubGal80^ts^ flies were raised in a Torrey Pines IN35 programable incubator where the temperature was automatically cycled from 18° to 25° twice per week for either 12 or 24 hours.

### RNA-Seq Analysis

RNA-seq was aligned against BDGP6.22 (Ensembl version 97) using STAR v2.7.1a (*66*), and quantified using RSEM v1.3.1 (*67*). Reads mapping to genes annotated as rRNA, snoRNA, or snRNA were removed. Genes which had less than 10 reads mapping on average across all samples were also removed. A differential expression analysis was performed using DESeq2 (*68*). The likelihood ratio test (LRT) was used to identify any genes that show change in expression across the different conditions. Pairwise comparisons were performed using a Wald test, with independent filtering. To control for false positives due to multiple comparisons in the genome-wide differential expression analysis, the false discovery rate (FDR) was computed using the Benjamini–Hochberg procedure. The gene level counts were transformed using a regularized log transformation, converted to z-scores, and clustered using partitioning around medoids (PAM), using correlation distance as the distance metric. Gene ontology (GO) and KEGG pathway enrichments for each cluster were performed using EnrichR (*69-71*). Terms with an FDR < 10% were defined as significantly enriched.

### Mortality Analysis

All analysis of lifespan data and curve fitting was performed using the nls non-linear least square tools in the R programming language. Survival data was imported into R and a survival curve derived from Gompertz–Makeham mortality law was fitted according to (*41, 42*). Briefly, survival curves are the integral of (Eq. 1), that is: Survival at a given age can be expressed as exp(A*MRTD/ln(2)*(1-exp(ln(2)*age/MRTD)-B*age). The B term captures death of flies due to age-independent causes such as sticking to food or transfer injury. B was fixed empirically to a low estimate of 0.001 or 0.1% of the total cohort per day. The MRTD and A parameters were then fitted to the empirical survival data using the nls library functions in R. Confidence intervals for the A and MRTD and residual standard errors were generated as part of the non-linear fit. For statistical testing, two parameters were considered statistically significantly different if their 95% confidence intervals did not overlap. The hypothetical mortality and survival statistic for the combination treatments were generated by applying fold changes of both individual interventions to MRTD and A parameters for each separate intervention sequentially.

## FIGURE LEGENDS

**Supplementary Figure 1**: **(A)** Schematic of the Drosophila digestive system comprising enterocytes (ECs), enteroendocrine (EEs), enteroblasts (EBs) and intestinal stem cells (ISCs). ISCs are located externally, away from the lumen. They divide symmetrically to make more ISCs or asymmetrically to form EBs or transit amplifying cells that further differentiate into ECs and EEs. Model for the the two treatments, rejuvenating differentiated cells to more stem cell like cells through Yamanaka factor expression, and activating apoptosis of senescent cells through peptide-based interference in FoxO-p53 binding.

**Supplementary Figure 2**: **Lifespan study of constitutive OKSM, Sen and OKSM-Sen expression in fly guts**. (A) Expression of OKSM, Sen and OKSM-Sen in *escargot* expressing cells of guts of adult female flies resulted in significantly reduced lifespan as compared to control flies (esgGFP TdTom). (B) Induced expression of OKSM, Sen and OKSM-Sen (post-eclosion) in separated male and female flies using drug induced expression resulted in significantly increased lifespans for both male and female flies. A P-value of 0 reflects P < 1.0 * 10^−10^.

**Supplementary Figure 3**: **Exploratory Data Analysis (EDA) of gene expression changes in the *Drosophila* gut in Sen, OKSM and OKSM-Sen treatment**. (A) PCA plots of gene expression in ubiquitous expression experiments with armGal4; tubGal80^ts^ > UAS-TdTomato (WT), armGal4; tubGal80^ts^ > UAS-OKSM (OKSM), armGal4; tubGal80^ts^ > UAS-Sen (Sen) and armGal4; tubGal80^ts^ > UAS-Sen; UAS-OKSM (OKSM-Sen). (B) PCA plots of gene expression in stem cell only expression experiments with esgGal4; tubGal80^ts^ > UAS-TdTomato (WT), esgGal4; tubGal80^ts^ > UAS-OKSM (OKSM), esgGal4; tubGal80^ts^ > UAS-Sen (Sen) and esgGal4; tubGal80^ts^ > UAS-Sen; UAS-OKSM (OKSM-Sen). (C) Z-scores boxplots of ubiquitous expression experiments with armGal4 for each of the seven clusters (D) Z-scores boxplots of ISC-restricted expression model with esgGal4 for each of the seven clusters.

**Supplementary Table 1**: Differential expression analysis results for ubiquitous expression model.

**Supplementary Table 2**: Differential expression analysis results for ISC-restricted expression model.

**Supplementary Table 3:** Fitting parameters and statistics from Gompertz–Makeham mortality and survival analysis. Best fit value and 95% confidence interval are listed for each parameter. Mortality rate doubling time is given in days and rounded to one decimal place. Initial mortality parameter A is normalized to that of the relevant control cohort (WT flies subjected to the same induction condition). All A parameter values are expressed in percent of this control.

## Data availability

RNA-seq data from this study has been deposited to GEO (X).

## Code availability

All code necessary to recreate the results from the analysis presented is available from: https://github.com/harmstonlab/OKSM_Senolytic

## Author contributions

Conceptualization, N.S.T.; methodology, P.K., E.H.Z.C., A.O. and A.R.; software E.H.Z.C.; modelling, J.G.; investigation, P.K., E.H.Z.C., A.O. and A.R.; resources, N.H., J.G., and N.S.T.; writing—original draft preparation, N.S.T.; writing—review and editing, P.K., E.H.Z.C., N.H., A.R, A.O, J.G., and N.S.T.; funding acquisition, N.S.T. and N.H. All authors have read and agreed to the published version of the manuscript.

## Acknowledgements

This work was supported by Ministry of Education, AcRF grants IG19-SI102 and IG20-BG101to NST, National University of Singapore and Yale-NUS College (through Reimagine Research Grant IG20-RRSG-001) to NH.

## REFERENCES

1. M. D. W. Piper, L. Partridge, Drosophila as a model for ageing. Biochimica et Biophysica Acta (BBA) - Molecular Basis of Disease 1864, 2707–2717 (2018).

2. P. Kaur, H. J. Jin, J. B. Lusk, N. S. Tolwinski, Modeling the Role of Wnt Signaling in Human and Drosophila Stem Cells. Genes (Basel) 9, (2018).

3. L. Hayflick, P. S. Moorhead, The serial cultivation of human diploid cell strains. Exp Cell Res 25, 585–621 (1961).

4. M. Serrano, A. W. Lin, M. E. McCurrach, D. Beach, S. W. Lowe, Oncogenic ras provokes premature cell senescence associated with accumulation of p53 and p16INK4a. Cell 88, 593–602 (1997).

5. J. Krishnamurthy et al., Ink4a/Arf expression is a biomarker of aging. The Journal of clinical investigation 114, 1299–1307 (2004).

6. J. N. Justice et al., Senolytics in idiopathic pulmonary fibrosis: Results from a first-in-human, open-label, pilot study. EBioMedicine 40, 554–563 (2019).

7. K. Takahashi, S. Yamanaka, Induction of pluripotent stem cells from mouse embryonic and adult fibroblast cultures by defined factors. Cell 126, 663–676 (2006).

8. G.-H. Liu et al., Recapitulation of premature ageing with iPSCs from Hutchinson– Gilford progeria syndrome. Nature 472, 221–225 (2011).

9. M. Abad et al., Reprogramming in vivo produces teratomas and iPS cells with totipotency features. Nature 502, 340–345 (2013).

10. K. Ohnishi et al., Premature termination of reprogramming in vivo leads to cancer development through altered epigenetic regulation. Cell 156, 663–677 (2014).

11. A. Ocampo, J. C. Izpisua Belmonte, Stem cells. Holding your breath for longevity. Science 347, 1319–1320 (2015).

12. A. Ocampo, P. Reddy, J. C. I. Belmonte,Anti-Aging Strategies Based on Cellular Reprogramming. Trends Mol Med 22, 725–738 (2016).

13. A. Ocampo et al., In Vivo Amelioration of Age-Associated Hallmarks by Partial Reprogramming. Cell 167, 1719–1733 e1712 (2016).

14. W. Zhang, J. Qu, G. H. Liu, J. C. I. Belmonte, The ageing epigenome and its rejuvenation. Nat Rev Mol Cell Biol 21, 137–150 (2020).

15. P. Reddy, S. Memczak, J. C. Izpisua Belmonte,Unlocking Tissue Regenerative Potential by Epigenetic Reprogramming. Cell Stem Cell 28, 5–7 (2021).

16. Y. Lu et al., Reprogramming to recover youthful epigenetic information and restore vision. Nature 588, 124–129 (2020).

17. A. Roux et al., Partial reprogramming restores youthful gene expression through transient suppression of cell identity. bioRxiv, 2021.2005.2021.444556 (2021).

18. T. J. Sarkar et al., Transient non-integrative expression of nuclear reprogramming factors promotes multifaceted amelioration of aging in human cells. Nat Commun 11, 1545 (2020).

19. D. Gill et al., Multi-omic rejuvenation of human cells by maturation phase transient reprogramming. Elife 11, e71624 (2022).

20. D. J. Baker et al., Clearance of p16Ink4a-positive senescent cells delays ageing-associated disorders. Nature 479, 232–236 (2011).

21. D. J. Baker et al., Naturally occurring p16Ink4a-positive cells shorten healthy lifespan. Nature 530, 184–189 (2016).

22. M. P. Baar et al., Targeted Apoptosis of Senescent Cells Restores Tissue Homeostasis in Response to Chemotoxicity and Aging. Cell 169, 132–147 e116 (2017).

23. M. Xu et al., Senolytics improve physical function and increase lifespan in old age. Nat Med 24, 1246–1256 (2018).

24. L. J. Hickson et al., Senolytics decrease senescent cells in humans: Preliminary report from a clinical trial of Dasatinib plus Quercetin in individuals with diabetic kidney disease. EBioMedicine 47, 446–456 (2019).

25. E. Dolgin, Send in the senolytics. Nat Biotechnol 38, 1371–1377 (2020).

26. E. O. Wissler Gerdes, Y. Zhu, T. Tchkonia, J. L. Kirkland, Discovery, development, and future application of senolytics: theories and predictions. FEBS J 287, 2418–2427 (2020).

27. S. He, N. E. Sharpless, Senescence in health and disease. Cell 169, 1000–1011 (2017).

28. T. D. Admasu et al., Drug Synergy Slows Aging and Improves Healthspan through IGF and SREBP Lipid Signaling. Dev Cell 47, 67–79 e65 (2018).

29. K. Takahashi et al. (2007).

30. J. Yu et al., Induced pluripotent stem cell lines derived from human somatic cells. Science 318, 1917–1920 (2007).

31. C. A. Sommer et al., Induced pluripotent stem cell generation using a single lentiviral stem cell cassette. Stem cells 27, 543–549 (2009).

32. R. A. Rossello et al., Mammalian genes induce partially reprogrammed pluripotent stem cells in non-mammalian vertebrate and invertebrate species. Elife 2, e00036 (2013).

33. C. A. Micchelli, N. Perrimon, Evidence that stem cells reside in the adult Drosophila midgut epithelium. Nature 439, 475–479 (2006).

34. B. Ohlstein, A. Spradling, The adult Drosophila posterior midgut is maintained by pluripotent stem cells. Nature 439, 470–474 (2006).

35. H. Jasper, Intestinal Stem Cell Aging: Origins and Interventions. Annu Rev Physiol 82, 203–226 (2020).

36. P. Zhang, B. A. Edgar, Insect Gut Regeneration. Cold Spring Harb Perspect Biol 14, a040915 (2022).

37. S. Nagarkar-Jaiswal et al., A genetic toolkit for tagging intronic MiMIC containing genes. Elife 4, e08469 (2015).

38. C. Neophytou, C. Pitsouli, How Gut Microbes Nurture Intestinal Stem Cells: A Drosophila Perspective. Metabolites 12, 169 (2022).

39. T. Zhao, Y. Xu, p53 and stem cells: new developments and new concerns. Trends Cell Biol 20, 170–175 (2010).

40. D. Rogulja, K. D. Irvine, Regulation of cell proliferation by a morphogen gradient. Cell 123, 449–461 (2005).

41. B. Gompertz, XXIV. On the nature of the function expressive of the law of human mortality, and on a new mode of determining the value of life contingencies. In a letter to Francis Baily, Esq. FRS &c. Philosophical transactions of the Royal Society of London, 513–583 (1825).

42. A. Kowald, T. B. Kirkwood, Senolytics and the compression of late-life mortality. Experimental Gerontology 155, 111588 (2021).

43. M. Demaria, P. Y. Desprez, J. Campisi, M. C. Velarde, Cell autonomous and non-autonomous effects of senescent cells in the skin. Journal of Investigative Dermatology 135, 1722–1726 (2015).

44. D. A. Guertin, K. V. Guntur, G. W. Bell, C. C. Thoreen, D. M. Sabatini, Functional genomics identifies TOR-regulated genes that control growth and division. Curr Biol 16, 958–970 (2006).

45. S. Haller et al., mTORC1 Activation during Repeated Regeneration Impairs Somatic Stem Cell Maintenance. Cell Stem Cell 21, 806-818.e805 (2017).

46. P. F. da Silva et al., The bystander effect contributes to the accumulation of senescent cells in vivo. Aging cell 18, e12848 (2019).

47. G. Nelson et al., A senescent cell bystander effect: Senescence-induced senescence. Aging cell 11, 345–349 (2012).

48. F. Zhou, A. Rasmussen, S. Lee, H. Agaisse, The UPD3 cytokine couples environmental challenge and intestinal stem cell division through modulation of JAK/STAT signaling in the stem cell microenvironment. Developmental biology 373, 383–393 (2013).

49. M. Xu, T. Tchkonia, J. L. Kirkland, Perspective: Targeting the JAK/STAT pathway to fight age-related dysfunction. Pharmacol Res 111, 152–154 (2016).

50. P. Houtz et al., Hippo, TGF-β, and Src-MAPK pathways regulate transcription of the upd3 cytokine in Drosophila enterocytes upon bacterial infection. PLoS genetics 13, e1007091 (2017).

51. K. Tominaga, H. I. Suzuki, TGF-β signaling in cellular senescence and aging-related pathology. International journal of molecular sciences 20, 5002 (2019).

52. A. Gibaja et al., TGFβ2-induced senescence during early inner ear development. Scientific reports 9, 1–13 (2019).

53. A. Rallis et al., Hedgehog Signaling Modulates Glial Proteostasis and Lifespan. Cell Rep 30, 2627–2643 e2625 (2020).

54. P. J. Ross et al., Human-induced pluripotent stem cells produced under xeno-free conditions. Stem Cells Dev 19, 1221–1229 (2010).

55. J. Bischof et al., A versatile platform for creating a comprehensive UAS-ORFeome library in Drosophila. Development 140, 2434–2442 (2013).

56. J. Bischof, R. K. Maeda, M. Hediger, F. Karch, K. Basler, An optimized transgenesis system for Drosophila using germ-line-specific φC31 integrases. Proceedings of the National Academy of Sciences 104, 3312–3317 (2007).

57. A. C. Groth, M. Fish, R. Nusse, M. P. Calos, Construction of transgenic Drosophila by using the site-specific integrase from phage φC31. Genetics 166, 1775–1782 (2004).

58. A. H. Brand, N. Perrimon, Targeted gene expression as a means of altering cell fates and generating dominant phenotypes. development 118, 401–415 (1993).

59. S. E. McGuire, P. T. Le, A. J. Osborn, K. Matsumoto, R. L. Davis, Spatiotemporal rescue of memory dysfunction in Drosophila. Science 302, 1765–1768 (2003).

60. K. J. Venken et al., MiMIC: a highly versatile transposon insertion resource for engineering Drosophila melanogaster genes. Nat Methods 8, 737–743 (2011).

61. S. Nagarkar-Jaiswal et al., A library of MiMICs allows tagging of genes and reversible, spatial and temporal knockdown of proteins in Drosophila. Elife 4, e05338 (2015).

62. N. A. Kaplan, P. F. Colosimo, X. Liu, N. S. Tolwinski, Complex interactions between GSK3 and aPKC in Drosophila embryonic epithelial morphogenesis. PLoS One 6, e18616 (2011).

63. P. Kaur et al., Wnt Signaling Rescues Amyloid Beta-Induced Gut Stem Cell Loss. Cells 11, (2022).

64. S. Han et al., OASIS 2: online application for survival analysis 2 with features for the analysis of maximal lifespan and healthspan in aging research. Oncotarget 7: 56147–56152. Trends Cell Biol 26, 565–568 (2016).

65. S. K. Han et al., OASIS 2: online application for survival analysis 2 with features for the analysis of maximal lifespan and healthspan in aging research. Oncotarget 7, 56147–56152 (2016).

66. A. Dobin et al., STAR: ultrafast universal RNA-seq aligner. Bioinformatics 29, 15–21 (2013).

67. B. Li, C. N. Dewey, RSEM: accurate transcript quantification from RNA-Seq data with or without a reference genome. BMC bioinformatics 12, 1–16 (2011).

68. M. I. Love, W. Huber, S. Anders, Moderated estimation of fold change and dispersion for RNA-seq data with DESeq2. Genome Biol 15, 550 (2014).

69. E. Y. Chen et al., Enrichr: interactive and collaborative HTML5 gene list enrichment analysis tool. BMC bioinformatics 14, 1–14 (2013).

70. M. V. Kuleshov et al., Enrichr: a comprehensive gene set enrichment analysis web server 2016 update. Nucleic acids research 44, W90–W97 (2016).

71. Z. Xie et al., Gene set knowledge discovery with Enrichr. Current protocols 1, e90 (2021).

