## Supplemental Data 1 for "Combining Stem Cell Rejuvenation and Senescence Targeting to Synergistically Extend Lifespan"

A. Drosophila Gastric System Schematic

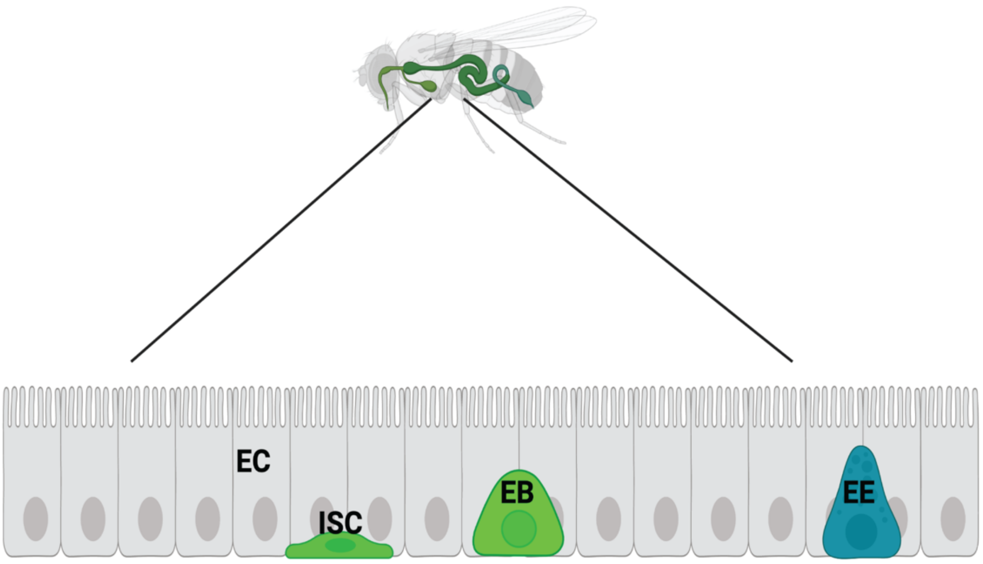


B. Two Interventions: Cell Rejuvenation and Senescent Cell Removal

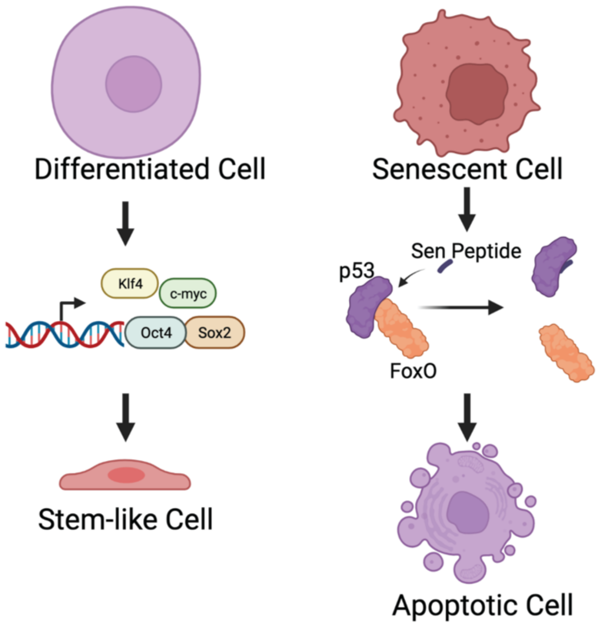


**Supplementary Figure 1**: **(A)** Schematic of the Drosophila digestive system comprising enterocytes (ECs), enteroendocrine (EEs), enteroblasts (EBs) and intestinal stem cells (ISCs). ISCs are located externally, away from the lumen. They divide symmetrically to make more ISCs or asymmetrically to form EBs or transit amplifying cells that further differentiate into ECs and EEs. **(B)** Model for the the two treatments, rejuvenating differentiated cells to more stem cell like cells through Yamanaka factor expression, and activating apoptosis of senescent cells through peptide-based interference in FoxO-p53 binding.

A. Constant ISC Only Expression Lifespan

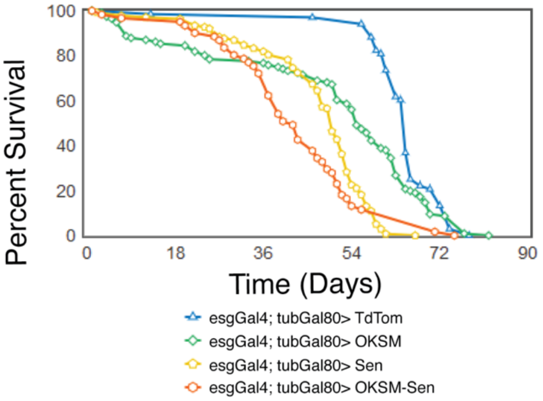


| Condition | Number of samples | Mean lifespan | | Max Lifespan (90th percentile) | |
|---|---|---|---|---|---|
| | | Age in days | p-value versus TdTom (Log-rank test) | Age in days | p-value versus TdTom (Fisher's exact test) |
| esgGAL4; UAS-GFP; TubGal80 TdTom | 68 | 64.41 | - | 74.00 | - |
| esgGAL4; UAS-GFP; TubGal80 OKSM | 116 | 49.66 | 0.003 (**) | 70.00 | 0.215 |
| esgGAL4; UAS-GFP; TubGal80 Sen | 138 | 46.60 | 0.000 (**) | 59.00 | 0.000 (**) |
| esgGAL4; UAS-GFP; TubGal80 OKSM-Sen | 61 | 42.57 | 0.000 (**) | 71.00 | 0.019 (*) |

B. Weekly Cycling Expression Lifespan RU486

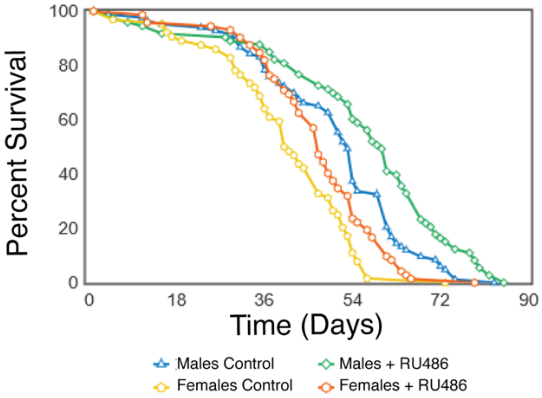


| Gender | Condition | Number of samples | Mean lifespan | | Max Lifespan (90th percentile) | |
|---|---|---|---|---|---|---|
| | | | Age in days | p-value, Ethanol versus RU486 (Log-rank test) | Age in days | p-value, Ethanol versus RU486 (Fisher's exact test) |
| Actin-Switch GFP; UAS-OKSM Males | Ethanol Control | 83 | 50.04 | 0.006 (**) | 68.00 | 0.054 |
| | RU486 | 73 | 55.86 | | 79.00 | |
| Actin-Switch GFP; UAS-OKSM Females | Ethanol Control | 64 | 40.41 | 0.018 (*) | 55.00 | 0.003 (**) |
| | RU486 | 72 | 46.68 | | 61.00 | |

**Supplementary Figure 2**: **Lifespan study of constitutive OKSM, Sen and OKSM-Sen expression in fly guts**. (A) Expression of OKSM, Sen and OKSM-Sen in *escargot* expressing cells of guts of adult female flies resulted in significantly reduced lifespan as compared to control flies (esgGFP TdTom). (B) Induced expression of OKSM, Sen and OKSM-Sen (post-eclosion) in separated male and female flies using drug induced expression resulted in significantly increased lifespans for both male and female flies. A P-value of 0 reflects $P < 1.0 * 10^{-10}$.

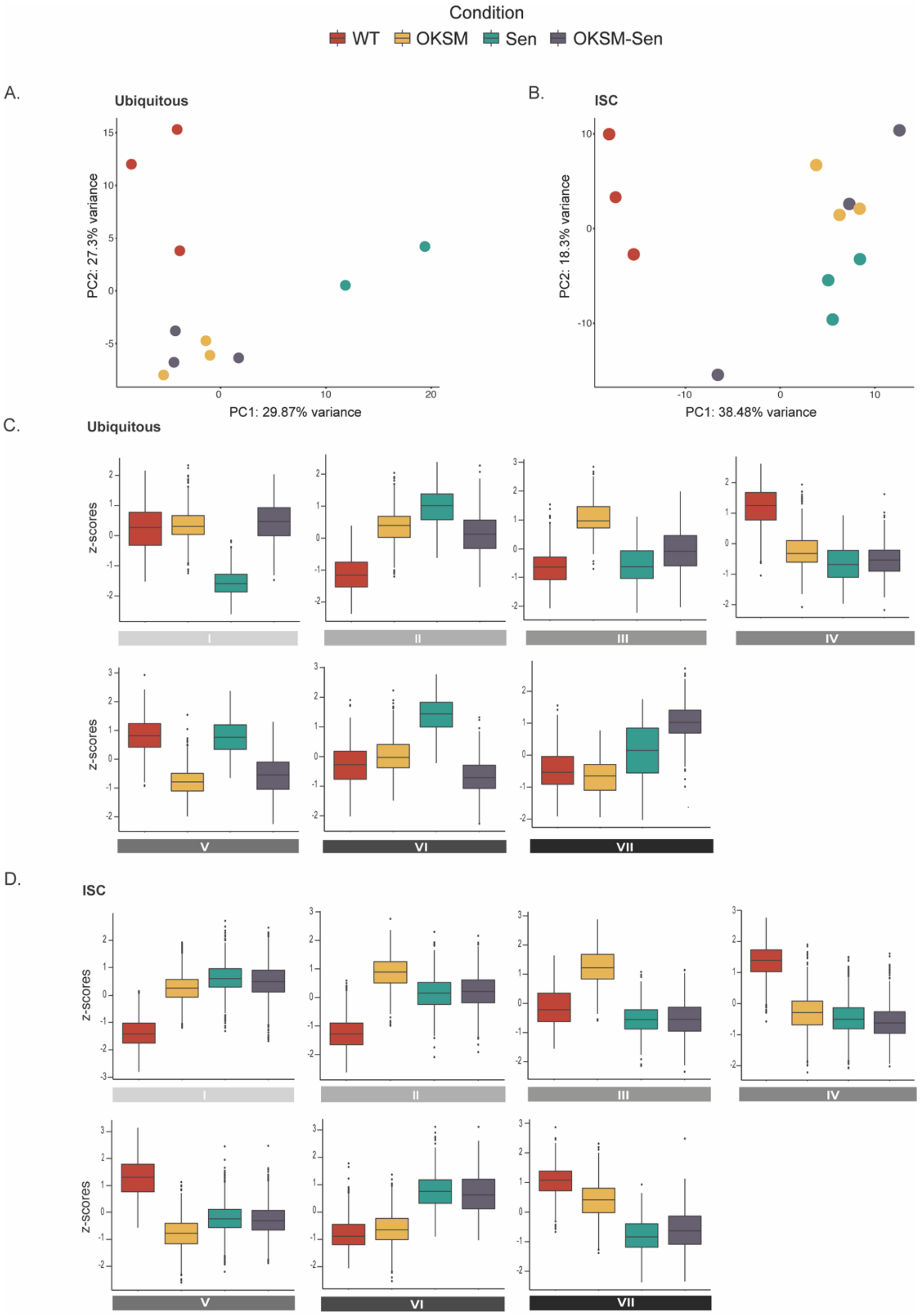


**Supplementary Figure 3**: **Exploratory Data Analysis (EDA) of gene expression changes in the *Drosophila* gut in Sen, OKSM and OKSM-Sen treatment.** (A) PCA plots of gene expression in ubiquitous expression experiments with armGal4; tubGal80$^{ts}$ > UAS-TdTomato (WT), armGal4; tubGal80$^{ts}$ > UAS-OKSM (OKSM), armGal4; tubGal80$^{ts}$ > UAS-Sen (Sen) and armGal4; tubGal80$^{ts}$ > UAS-Sen; UAS-OKSM (OKSM-Sen). (B) PCA plots of gene expression in stem cell only expression experiments with esgGal4; tubGal80$^{ts}$ > UAS-TdTomato (WT), esgGal4; tubGal80$^{ts}$ > UAS-OKSM (OKSM), esgGal4; tubGal80$^{ts}$ > UAS-Sen (Sen) and esgGal4; tubGal80$^{ts}$ > UAS-Sen; UAS-OKSM (OKSM-Sen). (C) Z-scores boxplots of ubiquitous expression experiments with armGal4 for each of the seven clusters (D) Z-scores boxplots of ISC-restricted expression model with esgGal4 for each of the seven clusters.

**Supplementary Table 1**: Differential expression analysis results for ubiquitous expression model.

**Supplementary Table 2**: Differential expression analysis results for ISC-restricted expression model.

**Supplementary Table 3:** Fitting parameters and statistics from Gompertz–Makeham mortality and survival analysis. Best fit value and 95% confidence interval are listed for each parameter. Mortality rate doubling time is given in days and rounded to one decimal place. Initial mortality parameter A is normalized to that of the relevant control cohort (WT flies subjected to the same induction condition). All A parameter values are expressed in percent of this control.
